# Detecting Anomalous Proteins Using Deep Representations

**DOI:** 10.1101/2023.04.03.535457

**Authors:** Tomer Michael-Pitschaze, Niv Cohen, Dan Ofer, Yedid Hoshen, Michal Linial

## Abstract

Many advances in biomedicine can be attributed to identifying unusual proteins and genes. Many of these proteins’ unique properties were discovered by manual inspection, which is becoming infeasible at the scale of modern protein datasets. Here, we propose to tackle this challenge using anomaly detection methods that automatically identify unexpected properties. We adopt a state-of-the-art anomaly detection paradigm from computer vision, to highlight unusual proteins. We generate meaningful representations without labeled inputs, using pretrained deep neural network models. We apply these protein language models (pLM) to detect anomalies in function, phylogenetic families, and segmentation tasks. We compute protein anomaly scores to highlight human prion-like proteins, distinguish viral proteins from their host proteome, and mark non-classical ion/metal binding proteins and enzymes. Other tasks concern segmentation of protein sequences into folded and unstructured regions. We provide candidates for rare functionality (e.g., prion proteins). Additionally, we show the anomaly score is useful in 3D folding-related segmentation. Our novel method shows improved performance over strong baselines and has objectively high performance across a variety of tasks. We conclude that the combination of pLM and anomaly detection techniques is a valid method for discovering a range of global and local protein characteristics.

## 1. INTRODUCTION

The unprecedented growth in quality and quantity of molecular data (e.g., genomes, transcriptomes) in recent years raises the need for a systematic approach for functional annotation of new protein sequences (Friedberg, 2006). The consistent increase in the success of automatic functional prediction was mostly attributed to the expansion in the number and variety of solved structures and the overall increase in databases (Radivojac, et al., 2013). The Gene Ontology (GO) framework is used as a gold standard for the assignment of biochemical function, biological process, and cellular localization terms to proteins (Martin, et al., 2004). Protein function is intrinsically complex and poorly defined. It is mostly indirectly studied through evolutionary conservation (e.g., protein homologous families and 3D structure (Ofran, et al., 2005). However, proteins carry numerous cellular functions that are context dependent. Examples are protein-protein interactions, cell signaling, and the regulatory network. For the genomic-based collection of UniProtKB/TrEMBL (230 M sequences, Release 4/2022), less than 1% have experimental evidence, and the majority of the database includes predicted proteins with no supporting evidence (Ouzounis and Karp, 2002). With such constraints, direct experiments are the best way to determine high accuracy in functional prediction.

The assignment of functions to sequences is a challenging task. Even with known homologs, inheritance transitivity is a source of inaccuracy and ambiguity in multi-domain protein functions (Green and Karp, 2005). Annotation efforts usually fail when function is rare and represented by orphan sequences (Tautz and Domazet-Lošo, 2011). Occasionally, protein functions that were not previously observed are reported, which suggests that unique and unexpected functions exist and that methodologies for their systematic findings are needed. Examples are short peptides in humans that resemble snail cone toxins (Kaplan, et al., 2007), heat resistant *hero* proteins (Tsuboyama, et al., 2020), prion proteins that drive pathological aggregations (Halfmann, et al., 2010), intrinsically disordered proteins that involve phase separation (Uversky and Dunker, 2010) and more. Distinguishing alterations in protein function due to mistakes in translation (Drummond and Wilke, 2009), and developing effective screening methods to identify novel functions is of ultimate importance to the field of protein design and engineering (Ufarte, et al., 2015).

Deep learning approaches have led to protein fold prediction with extremely high accuracy, as implemented in AlphaFold. It also allowed populating the unstructured space of protein sequences with high confidence structures (Varadi, et al., 2022), with 36% of all amino acids in the human proteome predicted with high confidence (Tunyasuvunakool, et al., 2021). However, in instances where only a few homologs exist, or there is low divergence, AlphaFold predictions are of lower quality. Importantly, mapping a fold to its function is not always evident, as the same fold may account for a large number of unrelated functions (Orengo, et al., 1994). In addition, training models from sequence (Wan and Jones, 2020) using NLP methodologies (Khurana, et al., 2022; Ofer, et al., 2021), led to breakthroughs in protein function inference (e.g., ProteinBERT, ProtTrans, ESM) and more (Brandes, et al., 2022; Chowdhury, et al., 2022; Elnaggar, et al., 2022; Rives, et al., 2021).

Most deep learning approaches, both for proteins and other data modalities, heavily rely on manually annotated samples. However, often the most exciting research tasks require discovering new phenomena rather than distinguishing between known data classes. Breaking from previous research, we explore a new setting for identifying novel, previously unknown protein types. As such protein types are unknown and unexpected, we do not assume that any annotated examples of such proteins are provided to us. To detect such novel samples automatically, we must rely on the ability to distinguish between samples similar to the training data (‘*normal data*’) and novel data types (‘*anomalies*’). For example, in the simple case of tabular data, it was shown that *density estimation* of samples as vectors is a strong approach (Fischer, et al., 2008). Namely, a sample is deemed anomalous if its features are far away from any normal samples, so it is likely not to have come from the same distribution. With other data modalities, a key step for anomaly detection is to map sparse samples to a relatively dense embedding space (Ruff, et al., 2021). After mapping each sample to a descriptive vector, a density estimation approach can be applied to various data modalities, including images (Reiss, et al., 2021), time-series (Hoshen, 2022), tabular healthcare data (Cohen, et al., 2021; Gu, et al., 2019).

In this study, we aim to establish a computational method for anomaly detection in proteins at a genomic scale. The methods we adapt were originally developed and successfully applied in the domain of computer vision. We tune the method to seek novelty within different subgroups of the protein sequence space. For this goal, we introduce protein function annotation terms from Gene Ontology (GO) and UniProtKB keywords as ground truth. We also discuss functions that describe structural segments within protein sequences. We rely on the notion that proteins with the same 3D fold might share only minimal sequence similarity (Marks, et al., 2012; Webb and Sali, 2016). We present a method that highlights novel functions and discuss the predictive power needed to identify novel functions within the protein sequence database.

## 2. METHODS

We present a method for detecting anomalous proteins. Crucially, our method does not assume that we are able to characterize the anomalous proteins, as unusual and interesting proteins are often unexpected. We adapted recent breakthroughs in image anomaly detection (e.g., DN2 (Bergman, et al., 2020), SPADE (Cohen and Hoshen, 2020), PANDA (Reiss, et al., 2021)) and applied them to the amino acid sequence of proteins. Our method tackles different categories in protein function detection: full-length proteins (i.e., whether the whole protein is an enzyme or not) and at the local residue level, denoted as *protein anomaly segmentation*. Our approach for anomaly detection consists of two stages: deep protein feature extraction (section **2.1**) and anomaly scoring (section **2.2**). In section **2.3,** we describe our embedding method for full-length protein sequences.

### 2.1 Feature extraction

Anomaly detection methods require powerful representations of the data. We desire representations that reflect the biological semantic similarity between proteins. While proteins are encoded as amino acid sequences, which are simply represented as a sequence of characters, this does not explicitly hold information about their structure, roles and interactions within the protein. Many approaches have been developed for protein representation in the field of protein structure, protein-protein interactions (PPI) and function in general (Ben-Hur, et al., 2008; Brandes, et al., 2022).

In this work, we choose to represent protein sequences by deep neural network (NN) encoders. The encoders are first pretrained on huge protein datasets, to solve a bidirectional language modeling task (Devlin, et al., 2018; Rives, et al., 2021). Specifically, the model is trained on a subsequence of amino acids in the protein, where some have been replaced with masked out tokens and its task is to predict these “hidden” tokens. This paradigm has two main advantages: (i) it can exploit a massive, unlabelled protein dataset; (ii) a success on this task implicitly requires a high-level understanding of the protein, its function and structure. We assume that such powerful encoders have already been trained and provided to us. This assumption is realistic and based on the use of protein language models (pLM) that are readily available and were found highly effective for different downstream tasks. For example, ESM, ProtTrans, ProteinBERT (Brandes, et al., 2022; Elnaggar, et al., 2022; Rives, et al., 2021). Following common practice, we use the penultimate layer of the pretrained encoders as our representation. We denote the activation map for protein *P* as *ψ*( *P* ).The model provides a vector representation for each amino acid. We denote the representation of the i-th amino acid as *F_i_* = *Ψ*(*P*)_*i*_.

### 2.2 Anomaly scoring

We use a density-based anomaly scoring rule. The motivation is that normal proteins are common, and we often find multiple examples for each normal protein pattern. Conversely, anomalous proteins are anticipated to be rare, especially in model organisms that have been extensively studied. We will not expect to find many examples of each anomalous protein pattern. We therefore use *k* nearest neighbors (*k*NN) to measure the anomaly score of a particular protein or residue. Consider, for example, the representation of residue *i* in protein *P*, *Ψ*(*P*)_*i*_. We compute its distance from each residue embedding in all proteins in the training set. The anomaly score of our target protein residue *s*(*P,i*) is computed as the average of the distance to the *k* nearest residues. Let us denote the *k* nearest residues from the normal train data of our target residue *Ψ*(*P*)_*i*_ as *N_k_*(*Ψ*(*P*)^*i*^):

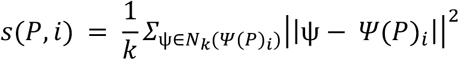

### 2.3 Whole sequence anomaly detection

Proteins can be anomalous either due to an anomalous local region, or due to a protein-wide anomalous property. To detect anomalies in entire proteins, one might consider using the anomaly score of the most anomalous residue within the entire protein. Although this method sometimes achieves strong results, it often fails. The reason is that residue embeddings are relatively local, while anomalous properties might be global. We propose whole-protein anomaly scoring, which considers both local and global patterns. One simple way to account for all the protein positions is to represent an entire protein as the mean of each of its residue embeddings. The mean embedding is similar to average pooling, a standard way of summarizing a sequence of features. To detect anomalies using the mean embedding, we look for proteins whose mean embedding is far from the mean embedding of any protein in the normal train set. This is computed using the Euclidean distance to its K nearest neighbors. Such proteins found in low density areas are likely to come from a different distribution than normal ones.

Using a single mean embedding for the entire protein may not provide a sufficient description of the local variations within the protein. An alternative way to summarize features, which can work better, is to use set features to represent the protein as a set of its segments. We adapt the method of Tzachor and Hoshen (Tzachor and Hoshen, 2023) in using set features to represent an entire sequence (Supplementary **Text S1**). Differently from their work, we operate on deep representations rather than on the raw data, as they are far more informative for proteins. We score a protein as normal or anomalous using nearest neighbors, with the Euclidean distance between the set features. We denote our whole-protein score (by either method) for protein *P* as *s*(*P*). All details on the implementation of our methodology and robustness to impurities in the training sets are in Supplementary **Text S1.**

## 3. RESULTS

### 3.1. Detecting contaminating viral proteins in human proteome

We evaluate the effectiveness of anomaly detection methods for discovering unknown protein types. In our experimental protocol, we provide training and test sets. The training set consists of normal proteins only, and the test set consists of normal and anomalous proteins. We train our method based only on the normal training data, and use it to compute anomaly scores for each test set protein. Our evaluation includes multiple anomaly types according to the different datasets used.

We tested the ability of the anomaly detection method to identify viral sequence contamination, specifically by human-infecting viruses. The task of identifying pathogenic viral sequences with respect to their hosts is of clinical relevance. We tested the ability of the method to detect viral proteins with respect to the host human proteome. A dataset was compiled from the curated SwissProt database. For this task, we further filtered the viruses to keep only those that are associated with humans as hosts. Out of 35K proteins, 27K remained following filtering by the host. See details in Supplementary **Text S1** and **Table S1**.

The accuracy of an anomaly detector is dependent on the desired tradeoff between false positive (FP) and false negative (FN) detections. In order to specify the desired tradeoff, one often chooses a threshold such that all samples with higher anomaly scores are considered anomalous. However, methods can have inconsistent ranking depending on the choice of threshold. For threshold-independent evaluation, most anomaly detection papers use the ROC-AUC metric which averages the true-positive rate for all possible false-negative rates (determined by different choices of threshold). An interpretation of this metric is that given a random normal sample *x_norm_* and an anomalous sample *x_anom_* (both from the test set) the ROC-AUC is equal to the probability that *s*(*x_anom_*) > *s*(*x_norm_*).

As further evidence for the efficacy of our approach, we computed the proportion of true anomalies out of the top M% proteins with the highest anomaly scores. Our anomaly score allowed us to prioritize candidate proteins with a far higher probability of being anomalous than randomly sampling the test set. **Table 1** summarizes the performance across multiple tasks. It is apparent that highly scored proteins have a far higher likelihood of being anomalous than randomly sampled ones.

**Table 1.** Frequency of anomalous samples for samples with high anomaly scores (% of the entire set).

| Set Features | Human / Virus | Enzyme | Extracellular / Intracellular | Ion / Metal binding | Multiple Interactions |
| --- | --- | --- | --- | --- | --- |
| Full Dataset | 75.00 | 21.55 | 17.33 | 20.34 | 45.81 |
| Test Set | 85.76 | 35.46 | 29.54 | 33.80 | 62.83 |
| Top 20% | 99.85 | 78.01 | 50.67 | 68.23 | 82.07 |
| Top 10% | 99.70 | 80.10 | 56.00 | 76.64 | 81.85 |
| Top 5% | 99.57 | 80.21 | 60.68 | 83.60 | 79.75 |
| Top 1% | 99.57 | 82.20 | 69.62 | 91.35 | 72.44 |

### 3.2. Detecting unknown protein types

We evaluate different datasets that cover diverse attributes of proteins, including biochemical functions, cellular localization, and protein interactions. We attempt to identify anomalies among enzymes (by their E.C. numbering system), proteins that belong to ion/metal binding proteins, and proteins with evidence for binding multiple proteins. For the latter, proteins were classified into two classes: those that interact with multiple other proteins and those that do not. We also modeled the subcellular localization of the proteins according to whether they were secreted/extracellular or located inside the cell. The set of secreted (i.e., their final location is in the extracellular space) proteins was evaluated versus any other cellular destination (referred to as “non-secreted”). For details on the prediction model and the database used, see Supplementary **Text S1**. Recall that a property such as being an ion-binding protein is based on the spatial arrangement of a small cluster of amino acids (i.e., local), while being involved in proteinprotein interaction has a more global context.

We compare our method against two baseline anomaly detection approaches. **Table 2** shows the results on these tasks. The ground truth for all the listed tasks is derived from SwissProt (unless stated otherwise). The *N-Gram* approach (Cavnar and Trenkle, 1994) relies on the raw amino-acid (encoded as one-hot vectors) counts rather than on deep representations of protein segments. As anticipated, it performs significantly worse, emphasizing the benefit of including contextualized embeddings from the pLMs (Ofer, et al., 2021). While the *Max of Segmentation* approach utilizes deep features, it underperforms the other approaches. The *mean embedding* method, which averages the features of protein segments, and the *set embedding* method, which uses set features, are the top performers on all datasets.

**Table 2.** Evaluation of anomaly detection methodologies.

| Dataset | Source | N-Gram | Max of Segmentation | Mean Embedding | Set Features |
| --- | --- | --- | --- | --- | --- |
| Human / Virus | SwissProt | 78.15 | 87.91 | 98.68 | 98.52 |
| Enzyme (E.C) | SwissProt | 67.78 | 74.33 | 84.54 | **85.32** |
| Extracellular / Intracellular | SwissProt | 60.87 | 57.61 | **74.91** | 70.59 |
| Ion / Metal binding | SwissProt | 66.37 | 66.58 | 78.69 | **79.93** |
| Multiple Interactions | SwissProt | 65.63 | 69.88 | 75.71 | **76.26** |
| Prions | UniRef90 | 78.05 | 29.23 | 80.59 | **85.76** |

### 3.3. Identifying novel prion-like proteins

To test the ability of our method to detect anomalies due to a global functional property, we tested our method on discovering prion-like characteristics and achieved 85.76% accuracy (ROC-AUC, *set embedding)*. The prion-anomaly detection dataset includes all non-fragment proteins from the UniRef90 database. We used UniRef90 proteins (i.e., representatives with <90% identity) in two manually annotated classes: known prion proteins that are labeled with the SwissProt molecular function keyword of Prion (KW-0640) versus all other (remaining) proteins. We focus on prions as these are rare with poorly defined biochemical properties. Prions undergo anomalous shifts in their 3D structure, which eventually leads to irreversible aggregation and physiopathology in vivo (Moore, et al., 2009).

Our unsupervised model was used to create a scored, ranked list of 60,000 prion predictions on a diverse sample of UniRef90 proteins. The top 100 most likely predictions are listed in Supplementary **Table S2**). We observed that the proteins with the highest anomaly scores are relatively short (mean 205 amino acids; length is <80 amino acids for 50% of the top list). The proteins are signified as having non-standard taxonomy sampling with over-representation of proteins from fungi and slime mold (14% each), bacteria and viruses (12% each). These are poorly studied proteins, with over a third of them being named “uncharacterized”. Many of these top scoring proteins have compositional bias (28%).

Our results support the notion that prion-like proteins have low sequence similarity to other proteins, low sequence complexity, and low confidence structure (**Fig. 2**). Most of the structures predicted as anomalous proteins are of low confidence (pLDDT orange/yellow color), and only very small fragments reach high confidence (dark blue). In many of these proteins, over-representation of specific amino acids is evident. For example, Q54VH6 and Q54QL5 from the Slime mold are composed of over 70% asparagine (N). Similarly, in Q54KT2, histidine (H) and proline (P) dominate the sequence. Extreme bias in the occurrence of amino acids signifies many of the identified prion-like proteins (Afsar Minhas, et al., 2017).

**Fig. 1.**
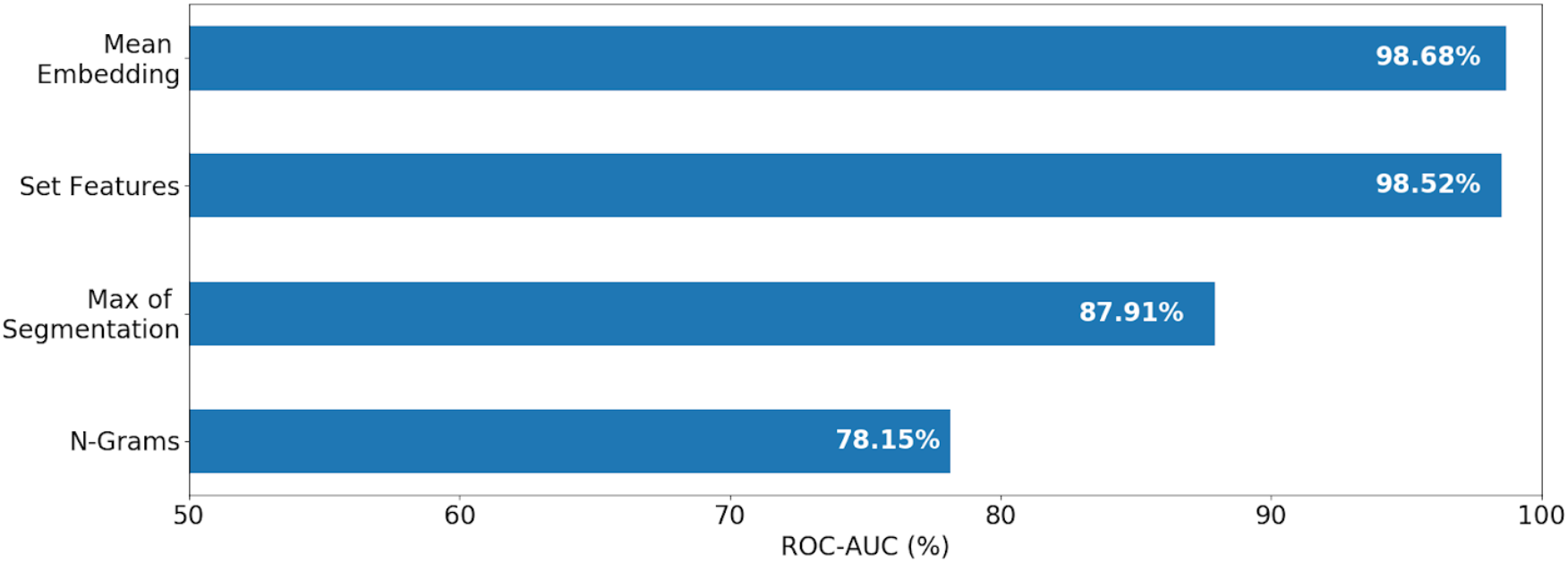
Anomaly detection accuracy for identifying human-specific viral proteins (% ROC-AUC) ROC-AUC performance in identifying contaminated viral proteins for the task of separating viral proteins from the host proteome. We only considered humans as viral hosts (and removed cases of broader virus-host tropism). Set and mean embedding methods achieved the best results.

**Fig. 2.**
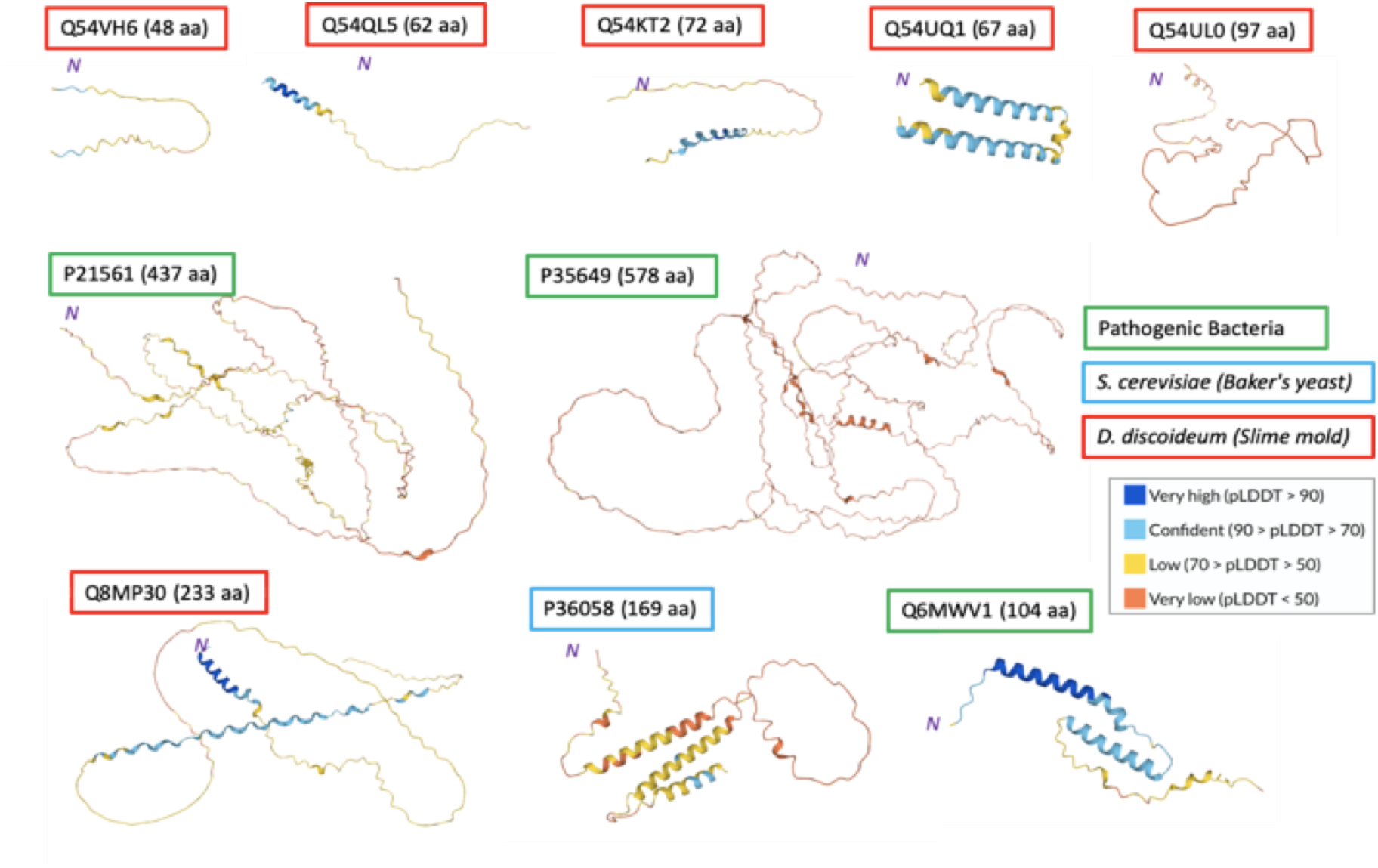
The 3D structures of the 10 proteins having the highest anomaly scores for the prion-like task. The AlphaFold2 structure predictions are shown and colored by the pLDDT confidence measure. The taxonomical groups are indicated by the color of the frames. Note that only small portions within these representative sequences are colored by high confidence (dark blue). Short proteins from the Slime mold dominated the anomalous prion-like sequences.

### 3.4. Correspondences between the anomaly score and structural-based segmentation

We further tested the ability of anomaly detection methods to assign local function and protein segmentation. To this end, we focused on 8,035 proteins and extracted residue-level anomaly scores for them. We analyzed a number of such cases and illustrate our findings for two representative proteins with respect to structural predictions by AlphaFold2 (Varadi, et al., 2022). Both proteins are characterized by long, unstructured segments. The UniProt Q96DN6 protein (1033 amino acids) is encoded by the gene MBD6 (Methyl-CpG-binding domain protein 6; **Fig. 3A)**. It binds to heterochromatin indirectly (without interacting with methylated or unmethylated DNA). In addition, it is recruited to sites of induced DNA damage and potentially acts in chromatin organization. While a detailed knowledge of its 3D is unavailable, AlphaFold2 predicts that the minimal positional error (dark green, **Fig. 3A,** middle) is limited to less than 100 amino acids at its N’-terminal domain. This region serves as an anchor site, with the rest of the 3D structure being of low confidence and a very large alignment positional error (**Fig. 3A,** bottom).

**Fig. 3.**
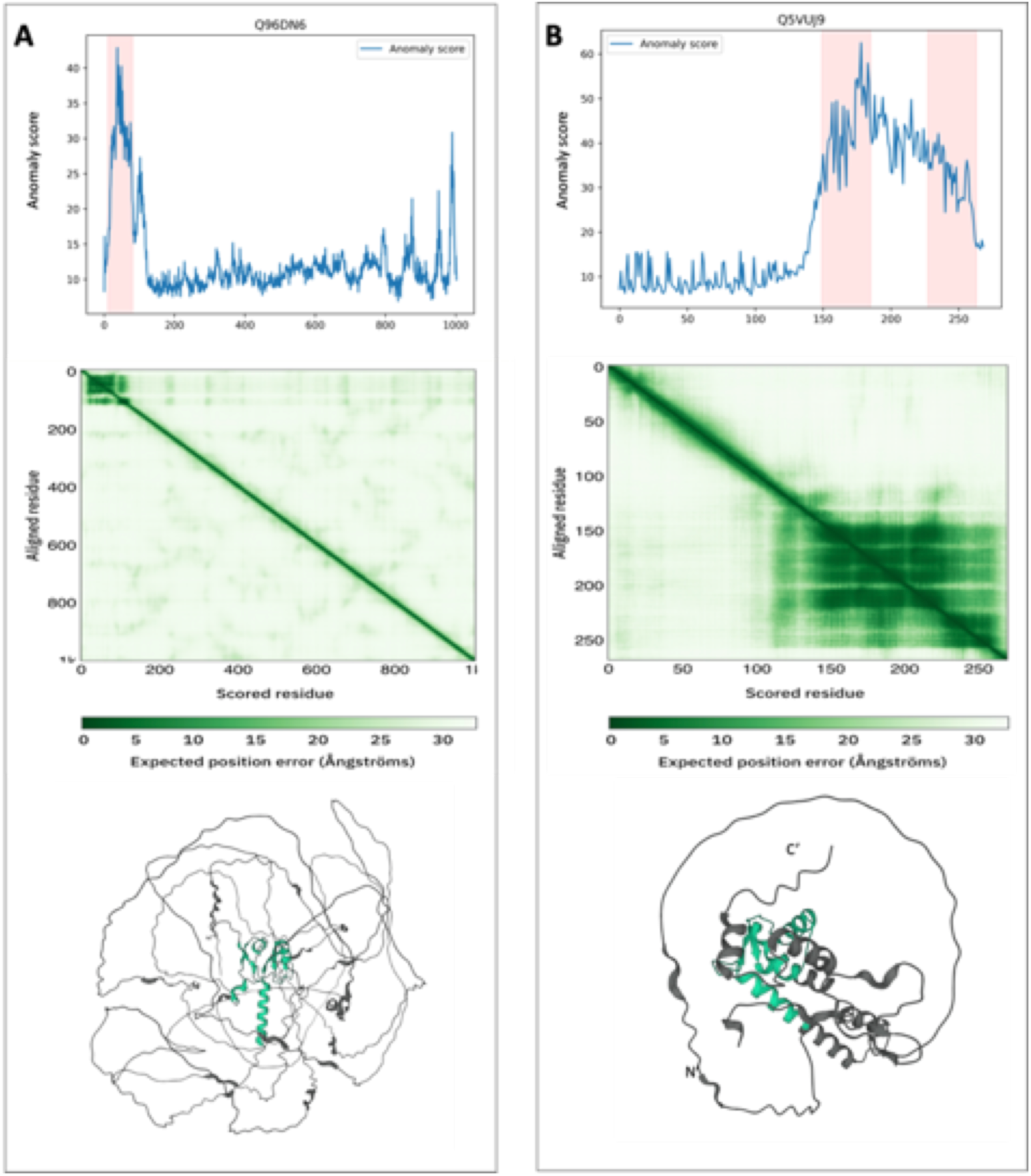
Residue-based anomaly detection protocol on represented proteins. **(A)** UniProtKB Q96DN6 (gene is MDB6), and **(B)** Q5VUJ9 (Dynein regulatory complex protein 8). Top: The plots show the anomaly score along the position of the sequence (x-axis). Note that the y-axis for the anomaly score is not identical in A and B. Pfam domains are colored pink. Middle: Predicted Aligned Error (PAE) plots. Specifically, the green color indicates expected distance error in Å. The color corresponds to the expected distance error in residue x’s position when the prediction and true structure are aligned on residue y. Bottom: AlphaFold2 predictions colored by the region with minimal PAE. Nate that the protein regions with maximal values of residue-based anomaly overlap the regions with minimal PAE values.

The second example of protein Q5VUJ9 (Dynein regulatory complex protein 8) has a similar trend. This protein (269 amino acids) regulates ciliary motility and the microtubule sliding in motile axonemes. The second half of the protein also acts as an anchor site for the extended low-confidence unstructured region (**Fig. 3B**). As shown in **Fig. 3B**, the unfolded segment that includes the first 200 amino-acids is poorly determined by AlphaFold2. Note that these unstructured long segments match the very low anomaly score. We conclude that the anomaly score detects non-classical proteins with large segments of unstructured regions, where high score highlights anchor regions (Fig. **3A** and Fig. **3B**, bottom). The anomaly score is seemingly less sensitive to domain boundaries. Although there is no direct information on the 3D structure or evolutionary conservation for many of the proteins marked as anomalous, the notion of folded region and non-structural regions is rediscovered by our methodology. Additional examples of the local anomaly score profiles are shown in Supplementary **Fig. S1**

## 4. DISCUSSION

Studying the source of anomalous proteins is relevant for understanding the source of functional novelty (Singh and Syrkin Wurtele, 2020). Identifying novel functions among the curated UniProtKB/SwissProt is a challenging task. Often it is restricted to proteins characterized by an accelerated evolutionary rate (i.e., under positive selection), enriched with mutations (i.e., polymorphic hotspot regions), or originating from a rapidly evolving phylogenetic lineage. Function annotations often rely on having homologues in model organisms. However, genuine anomalous proteins include orphan proteins, which are poorly annotated and overrepresented in less studied organisms (Hanson, et al., 2010). While the function of most proteins is uniquely defined, moonlighting proteins have multiple (often unrelated) functions. For example, the SMN’s (survival motor neuron) main role is in the biogenesis of small nuclear ribonucleoproteins (snRNPs). However, when expressed in axonal projections, it acts to control local translation (Sanchez, et al., 2013). Proteins may also alter their function according to their oligomerization state. The beta amyloid, which is the hallmark of Alzheimer’s disease, exhibits neurotoxicity when oligomerized. However, as a monomer, it actually acts to quench metal-inducible oxygen radicals, thereby inhibiting neurotoxicity (Zou, et al., 2002). Many of these unexpected examples were identified sporadically. We present a systematic method that can be used for identifying new candidates that are anomalous on a genomic scale.

Although our technical approach follows the leading paradigm for anomaly detection with deep features, future research may provide further improvements to our anomaly detection results. First, further improvement on deep protein embedding methods will directly improve the quality of our results. This is especially true, if we have some prior knowledge regarding some biological properties that may indicate anomalies. In this study, we applied a single approach to many different tasks (**Table 2**). However, it is possible that different embeddings may highlight different aspects of the proteins that will be useful for particular tasks. For example, proteins associated with functions that involve RNA or DNA must be quite long, often unstructured, and have abundant basic residues that are common in nucleic acid interactions from bacteria to humans. Incorporating such domain-specific knowledge can significantly improve performance.

Another possible direction for future improvements is using additional knowledge to adapt the used pretrained representation to better describe normal variation in the data and avoid false positive (FP) detection. Such knowledge may consist of statistical assumptions regarding the distribution of the normal data that can be used to adapt the representation. This was done in the cases of previous methods: DeepSVDD (Ruff, et al., 2018), PANDA (Reiss, et al., 2022), Mahalanobis (Rippel, et al., 2021), and OOD-no-labels (Cohen, et al., 2021). Prior knowledge may also come in the form of additional auxiliary features (Ofer, et al., 2021) or labels. While still not assuming labeled anomalies, the normal samples may have semantic labels. For example, class labels for protein function in the normal data may allow us to better adapt the representation to detect novel, unlabeled protein functions (Fort, et al., 2021; Hendrycks and Gimpel, 2016). Another option is that the user may label some attributes they wish to ignore. E.g., wishing not to detect known protein types in unseen organisms as anomalies. In such cases, we may provide organism labels for the data in order to make our representation agnostic to the source organism (Cohen, et al., 2022). A limitation in our search for anomalies stems from the generic protocol for protein annotations. For example, the automatic pipeline of genome annotation often overlooks short proteins (Hemm, et al., 2010; Linial, et al., 2017). Revisiting these short sequences revealed toxin-like function, novel antibiotics, and unexpected immunological cell recognition proteins (Kaplan, et al., 2007; Linial, et al., 2017).

We tested the ability of the anomaly detection method to identify viral sequence contamination (**Fig. 1**). From the results of this task, we were able to draw the following conclusions: (i) Human-host viruses were more likely to be detected as anomalies than viruses with a more general tropism. Viral proteins that were mistakenly classified as human proteins overlap with cases of mimicry (Rappoport and Linial, 2012). (ii) Latent viruses such as the herpes virus were misclassified as anomalies. Notably, latent viruses provide a real difficulty to the immune recognition system, where the separation between self to non-self is blurred. (iii) Retrovirus sequences are of viral origin, which along evolution became endogenous to the human genome. These are often misclassified as anomalies. Retroviral-like proteins are evolutionary remnants, and once integrated, their duplication is identical to any other human gene (Escalera-Zamudio and Greenwood, 2016).

In this study, we cover a wide range of protein functionalities, some of which are extremely rare (prions), while other functions are far more common (enzymes). Prion identification reached a high success rate. It may reflect the lack of contamination in the training, due to prion proteins’ rarity. Prions are of great interest from structural and medical perspectives. They act as infectious agents with devastating outcomes. From a biochemical point of view, the pathogenic protein may form non-reversible aggregates that lead to a chain reaction that infects benign prion proteins. Prion propagation is a common concept shared between mammals and fungi but has been poorly studied in other organisms (Tuite and Serio, 2010). Prion proteins may tilt the balance to accelerate the ‘infectious’ potential (Chakrabortee, et al., 2016). It was debated whether the infectivity capacity of prions is a true anomaly to our biological understanding (Malinovska, et al., 2013). Considering the unprecedented speed of determining protein sequences from poorly studied genomes, the unsupervised anomaly detection is an attractive approach for identifying functional novelty within the protein sequence database.

## Supplementary Materials

Supplementary Text S1 (Datasets, implementation, robustness to impurities)

Supplementary Fig. S1 (local anomaly scores-16 representative proteins).

Supplementary Table S1 (Human-viral set from SwissProt with annotations).

Supplementary Table S2 (100 top prion-like proteins and their FASTA sequences).

## Supplementary Text S1

### A. Dataset details

All datasets were derived from the curated SwissProt subset of the Uniprot database or the protein representatives of UniRef90. The datasets were based on the SwissProt reviewed collection of full-length proteins (removal of “fragment”). We created a combined dataset of human and virus proteins, including protein from viruses that affect ‘animals’. For the human/virus task, we further filtered the viruses to keep only those that defined their host as human (rather than animals). Altogether, the dataset was reduced from 35K proteins to 27K.

The Prions dataset included all non-fragment proteins from Uniref90 (i.e., including unreviewed proteins). Sequences were filtered to remove near duplicates based on the first N-terminal first 250 positions, with a reduced 18 letter amino acid alphabet (unified R and K, and I and V). Following this transformation, we were left with 26,981 proteins for the human/virus collection, out of 27,949. Ground truth labels were derived from SwissProt/UniProtKB unless stated otherwise. The tasks’ targets are defined as follows:

- Viral vs human proteins: Data are divided between viral (anomalous) and human (normal) proteins, according to origin proteome. Human endogenous retroviruses genes were also labeled as “human”.
- Enzymes: Data are divided between enzymes (anomalous) and non-enzymes (normal), according to the existence of an Enzyme Commission (E.C.) number.
- Ion/Metal binding: Proteins are divided into those bind ions/metals (anomalous) and proteins that do not (normal).
- Proteins that interact with multiple other proteins: Data are divided between proteins that interact with multiple other proteins (anomalous) and proteins that do not (normal). Interaction is defined from the “InteractsWith” attribute from SwissProt being greater than 1.
- Secreted subcellular localization: Proteins are divided according to their known subcellular locations: secreted (i.e., extracellular) proteins (anomalous) and non-secreted (normal). Proteins without known subcellular location are treated as negative (non-secreted) cases.
- Prion proteins: Classification of all UniRef90 proteins into two classes: manually reviewed, known Prion proteins (anomalous) with the SwissProt Prion (KW-0640) molecular function keyword and all others (normal).

Target class ratios were unbalanced as expected (binary tasks)

| Tasks | Minority class (%) |
| --- | --- |
| Prions (UR90) | 0.01% |
| Human/Virus | 25% |
| Enzyme | 43% |
| Extracellular/Intracellular | 36.8% |
| Ion/Metal binding | 18% |
| Multiple Interactions | 34% |

### B. Implementation Details

#### Preprocessing

Following ESM, all sequences were trimmed to the first 1022 amino acids.

#### Architecture

We used the ESM-1b (esm1b_t33_650M_UR50S) network, with weights provided by the authors of ESM.

#### Set features for protein sequences

We follow the method by (Tzachor and Hoshen, 2023). We use a window duration of 49 residues, 30 temporal dilations, 200 projections and 100-bin histograms. We did not use whitening.

#### Training set subsampling and cleaning

To improve efficiency and mitigate the potential presence of anomalies in the training set, we subsample the training set. In order to select the samples that best represent the normal data, our method picks the most *typical* samples. Concretely, we randomly select *n* (we used n=50,000) samples to be used as evaluators. For each of the remaining examples, we retrieve the K evaluators that are nearest to it, and average the distance to them. Finally, we select the *m* samples (we used m=50,000) that have the smallest K evaluator distance. For residue-level samples, nearby regions in the same protein are often embedded to very similar features. We ensure that evaluators are selected for a diverse set of proteins and locations. When fewer than n+m training samples are provided, we use random subsampling.

#### Runtime

Our method scores 3 proteins per second, using our unoptimized code and n=50,000. In naive implementations the runtime scales linearly with the size of the training set, but this is mitigated using our subsampling method.

#### Datasets

We used the UniProtKB/SwissProt section as a high-quality, curated database for most biological tasks. SwissProt (Release 2022_05, 570K proteins, also called “reviewed”) is a subset of manually reviewed, validated proteins. Among them 20K are from humans. We also used UniRef90 for the viral/human identification task. It is created by clustering the non-redundant UniRef100 sequences such that each cluster is composed of sequences that have >90% identity to the longest sequence in the cluster (i.e., the seed representative sequence). Furthermore, the proteins within a UniRef90 cluster will have >80% overlap in length with the seed sequence (Suzek, et al., 2015).

### C. Robustness to training set impurity

As anomalies are hard to detect, the training set may include anomalous samples. We test the robustness of our method to the existence of (unknown) anomalies in the training set (“impurity”). Figure below summarizes the impact of different levels of impurity in the training examining the enzymes dataset. First, we find that with a relatively small amount of data in the training dataset (up to 1% contamination), our results are not significantly degraded. When there is a larger degree of impurity in the training dataset (at the order of 10%), the degradation is more severe, but we are able to mitigate it using the selection of a representative subset described in Sec 2.4.

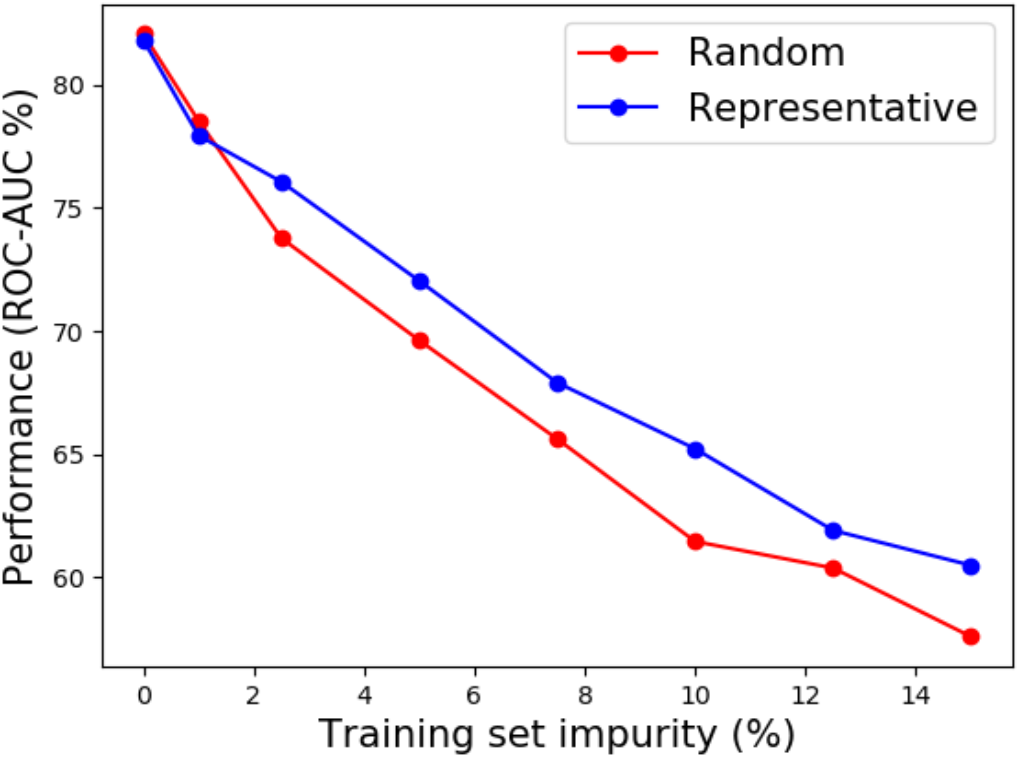

Anomaly detection accuracy for different impurity levels in the training set of the enzyme dataset. While very large impurity rates naturally reduce performance, our representative subsampling method improves performance over random subsampling.

---

**Figure S1.**
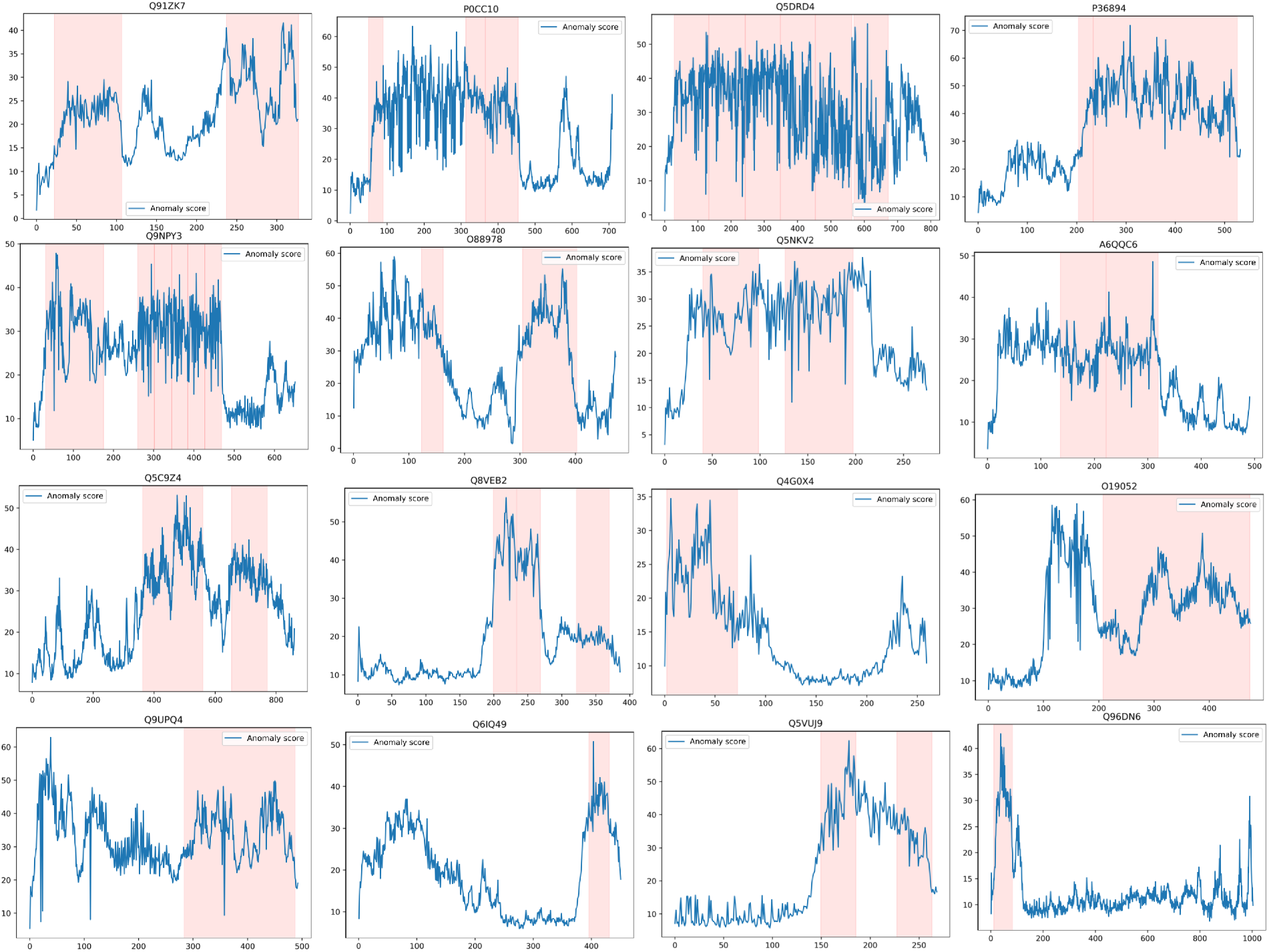
The plots show the anomaly score along the position of the sequence (x-axis). Note that the y-axis for the anomaly score is not identical in each panel. Pfam domains are colored pink.

## REFERENCES

Afsar Minhas, F.U.A., Ross, E.D. and Ben-Hur, A. Amino acid composition predicts prion activity. PLoS Comput Biol 2017;13(4):e1005465.

Ben-Hur, A., et al. Support vector machines and kernels for computational biology. PLoS computational biology 2008;4(10):e1000173.

Bergman, L., Cohen, N. and Hoshen, Y. Deep nearest neighbor anomaly detection. arXiv preprint arXiv:2002.10445 2020.

Brandes, N., et al. ProteinBERT: A universal deep-learning model of protein sequence and function. Bioinformatics 2022;38(8):2102–2110.

Cavnar, W.B. and Trenkle, J.M. N-gram-based text categorization. In, Proceedings of SDAIR-94, 3rd annual symposium on document analysis and information retrieval. Las Vegas, NV; 1994.

Chakrabortee, S., et al. Intrinsically disordered proteins drive emergence and inheritance of biological traits. Cell 2016;167(2):369–381. e312.

Chowdhury, R., et al. Single-sequence protein structure prediction using a language model and deep learning. Nature Biotechnology 2022;40(11):1617–1623.

Cohen, N., Abutbul, R. and Hoshen, Y. Out-of-Distribution Detection without Class Labels. arXiv preprint arXiv:2112.07662 2021.

Cohen, N. and Hoshen, Y. Sub-image anomaly detection with deep pyramid correspondences. arXiv preprint arXiv:2005.02357 2020.

Cohen, N., Kahana, J. and Hoshen, Y. Red PANDA: Disambiguating Anomaly Detection by Removing Nuisance Factors. arXiv preprint arXiv:2207.03478 2022.

Cohen, S., et al. ICU survival prediction incorporating test-time augmentation to improve the accuracy of ensemble-based models. IEEE Access 2021;9:91584–91592.

Devlin, J., et al. Bert: Pre-training of deep bidirectional transformers for language understanding. arXiv preprint arXiv:1810.04805 2018.

Drummond, D.A. and Wilke, C.O. The evolutionary consequences of erroneous protein synthesis. Nat Rev Genet 2009;10(10):715–724.

Elnaggar, A., et al. ProtTrans: Toward Understanding the Language of Life Through Self-Supervised Learning. IEEE Trans Pattern Anal Mach Intell 2022;44(10):7112–7127.

Escalera-Zamudio, M. and Greenwood, A.D. On the classification and evolution of endogenous retrovirus: human endogenous retroviruses may not be ‘human’after all. Apmis 2016;124(1-2):44–51.

Fischer, J., Mayer, C.E. and Söding, J. Prediction of protein functional residues from sequence by probability density estimation. Bioinformatics 2008;24(5):613–620.

Fort, S., Ren, J. and Lakshminarayanan, B. Exploring the limits of out-of-distribution detection. Advances in Neural Information Processing Systems 2021;34:7068–7081.

Friedberg, I. Automated protein function prediction—the genomic challenge. Briefings in bioinformatics 2006;7(3):225–242.

Green, M. and Karp, P. Genome annotation errors in pathway databases due to semantic ambiguity in partial EC numbers. Nucleic acids research 2005;33(13):4035–4039.

Gu, X., Akoglu, L. and Rinaldo, A. Statistical analysis of nearest neighbor methods for anomaly detection. Advances in Neural Information Processing Systems 2019;32.

Halfmann, R., Alberti, S. and Lindquist, S. Prions, protein homeostasis, and phenotypic diversity. Trends in cell biology 2010;20(3):125–133.

Hanson, A.D., et al. ‘Unknown’proteins and ‘orphan’enzymes: the missing half of the engineering parts list–and how to find it. Biochemical Journal 2010;425(1):1–11.

Hemm, M.R., et al. Small stress response proteins in Escherichia coli: proteins missed by classical proteomic studies. Journal of bacteriology 2010;192(1):46–58.

Hendrycks, D. and Gimpel, K. A baseline for detecting misclassified and out-of-distribution examples in neural networks. arXiv preprint arXiv:1610.02136 2016.

Hoshen, Y. Time Series Anomaly Detection by Cumulative Radon Features. arXiv preprint arXiv:2202.04067 2022.

Kaplan, N., Morpurgo, N. and Linial, M. Novel families of toxin-like peptides in insects and mammals: a computational approach. Journal of molecular biology 2007;369(2):553–566.

Khurana, D., et al. Natural language processing: State of the art, current trends and challenges. Multimedia tools and applications 2022:1–32.

Linial, M., Rappoport, N. and Ofer, D. Overlooked short toxin-like proteins: a shortcut to drug design. Toxins 2017;9(11):350.

Malinovska, L., Kroschwald, S. and Alberti, S. Protein disorder, prion propensities, and self-organizing macromolecular collectives. Biochimica et Biophysica Acta (BBA)-Proteins and Proteomics 2013;1834(5):918–931.

Marks, D.S., Hopf, T.A. and Sander, C. Protein structure prediction from sequence variation. Nature biotechnology 2012;30(11):1072–1080.

Martin, D., Berriman, M. and Barton, G.J. GOtcha: a new method for prediction of protein function assessed by the annotation of seven genomes. BMC bioinformatics 2004;5(1 ):1–17.

Moore, R.A., Taubner, L.M. and Priola, S.A. Prion protein misfolding and disease. Current opinion in structural biology 2009;19(1):14–22.

Ofer, D., Brandes, N. and Linial, M. The language of proteins: NLP, machine learning & protein sequences. Computational and Structural Biotechnology Journal 2021; 19:1750–1758.

Ofran, Y., et al. Beyond annotation transfer by homology: novel protein-function prediction methods to assist drug discovery. Drug discovery today 2005;10(21):1475–1482.

Orengo, C.A., Jones, D.T. and Thornton, J.M. Protein superfamilles and domain superfolds. Nature 1994;372(6507):631–634.

Ouzounis, C.A. and Karp, P.D. The past, present and future of genome-wide re-annotation. Genome Biol 2002;3(2):COMMENT2001.

Radivojac, P., et al. A large-scale evaluation of computational protein function prediction. Nature methods 2013;10(3):221–227.

Rappoport, N. and Linial, M. Viral proteins acquired from a host converge to simplified domain architectures. PLoS computational biology 2012;8(2):e1002364.

Reiss, T., et al. Panda: Adapting pretrained features for anomaly detection and segmentation. In, Proceedings of the IEEE/CVF Conference on Computer Vision and Pattern Recognition. 2021. p. 2806–2814.

Reiss, T., et al. Anomaly Detection Requires Better Representations. arXiv preprint arXiv:2210.10773 2022.

Rippel, O., Mertens, P. and Merhof, D. Modeling the distribution of normal data in pre-trained deep features for anomaly detection. In, 2020 25th International Conference on Pattern Recognition (ICPR). IEEE; 2021. p. 6726–6733.

Rives, A., et al. Biological structure and function emerge from scaling unsupervised learning to 250 million protein sequences. Proceedings of the National Academy of Sciences 2021;118(15):e2016239118.

Ruff, L., et al. A unifying review of deep and shallow anomaly detection. Proceedings of the IEEE 2021;109(5):756–795.

Ruff, L., et al. Deep one-class classification. In, International conference on machine learning. PMLR; 2018. p. 4393–4402.

Sanchez, G., et al. A novel function for the survival motoneuron protein as a translational regulator. Hum Mol Genet 2013;22(4):668–684.

Singh, U. and Syrkin Wurtele, E. How new genes are born. Elife 2020;9.

Suzek, B.E., et al. UniRef clusters: a comprehensive and scalable alternative for improving sequence similarity searches. Bioinformatics 2015;31(6):926–932.

Tautz, D. and Domazet Lošo, T. The evolutionary origin of orphan genes. Nature Reviews Genetics 2011;12(10):692–702.

Tsuboyama, K., et al. A widespread family of heat-resistant obscure (Hero) proteins protect against protein instability and aggregation. PLoS Biology 2020;18(3):e3000632.

Tuite, M.F. and Serio, T.R. The prion hypothesis: from biological anomaly to basic regulatory mechanism. Nat Rev Mol Cell Biol 2010;11 (12):823–833.

Tunyasuvunakool, K., et al. Highly accurate protein structure prediction for the human proteome. Nature 2021;596(7873):590–596.

Tzachor, I. and Hoshen, Y. Window Projection Features are All You Need for Time Series Anomaly Detection. ICLR 2023((under review)).

Ufarte, L., Potocki-Veronese, G. and Laville, E. Discovery of new protein families and functions: new challenges in functional metagenomics for biotechnologies and microbial ecology. Front Microbiol 2015;6:563.

Uversky, V.N. and Dunker, A.K. Understanding protein non-folding. Biochimica et Biophysica Acta (BBA)-Proteins and Proteomics 2010;1804(6):1231–1264.

Varadi, M., et al. AlphaFold Protein Structure Database: massively expanding the structural coverage of protein-sequence space with high-accuracy models. Nucleic acids research 2022;50(D1):D439–D444.

Wan, C. and Jones, D.T. Protein function prediction is improved by creating synthetic feature samples with generative adversarial networks. Nature Machine Intelligence 2020;2(9):540–550.

Webb, B. and Sali, A. Comparative protein structure modeling using MODELLER. Current protocols in bioinformatics 2016;54(1):5.6. 1–5.6. 37.

Zou, K., et al. A novel function of monomeric amyloid β-protein serving as an antioxidant molecule against metal-induced oxidative damage. Journal of Neuroscience 2002;22(12):4833–4841.

